# Immunopeptidomics reveals peptide antigens preferentially presented in psoriasis lesional skin of HLA-C*06:02 carriers

**DOI:** 10.64898/2026.01.24.701535

**Authors:** Bjørn Kromann Hansen, Shanzou Chung, Prithvi Raj Munday, Jingran Ye, Chen Li, Nathan P. Croft, Nicole A. Mifsud, Michael Bzorek, Varun Sharma, Aly Fayed, Graham Starkey, Rohit D’Costa, Claire L. Gordon, Morten Bahrt Haulrig, Thuvarahan Jegathees, Frances Burns, Johannes S. Kern, Lone Skov, Marianne Bengtson Løvendorf, Anthony W. Pumassmrcell, Beatrice Dyring-Andersen, Asolina Braun

## Abstract

**Background:** Human leukocyte antigen (HLA)-C*06:02 is a major genetic risk factor for psoriasis and understanding the HLA-C*06:02-presented peptide antigen repertoire (immunopeptidome) in the skin of patients is crucial for identifying autoantigens. Yet, no skin immunopeptidome data from patients stratified by their HLA-C*06:02 status exists.

**Objective:** We analysed biopsies from lesional and non-lesional skin of patients with psoriasis vulgaris (n=12), guttate psoriasis (n=8), or from skin of healthy controls (n=16).

**Methods:** HLA class I and class II peptide complexes were isolated by serial immunoprecipitation and HLA-bound peptides identified by liquid chromatography-tandem mass spectrometry. HLA-C*06:02 genotyping was performed by polymerase chain reaction.

**Results:** Over 99,000 non-redundant peptide ligands were identified across all samples. Substantially more HLA class I and class II peptides were detected in lesional psoriatic skin compared to matched non-lesional and healthy skin. Three peptides predicted to bind HLA-C*06:02, including MRASSFLIV from the known psoriasis marker peptidase inhibitor 3 (PI3), were identified in all lesions of HLA-C*06:02-positive patients but were rarely detected or absent in HLA-C*06:02-negative patient lesional skin and not detected at all in unaffected skin. Keratinocyte differentiation-associated protein (KRTDAP) was a notable source of lesion-specific HLA class II ligands contributing three out of six peptides detected in more than half of the lesional samples.

**Conclusion:** Active psoriatic lesions display an altered and expanded immunopeptidome compared to unaffected skin. We have identified numerous unreported, lesion-specific HLA-bound peptides and their source proteins. These findings offer insights into the pathobiology of psoriasis and provide a resource for future functional studies.

**CAPSULE SUMMARY:** A selection of immunopeptides is presented exclusively in lesional skin of HLA-C*06:02^+^ patients with psoriasis that may represent antigenic drivers of disease.

## Introduction

Psoriasis is a chronic, immune-mediated skin disease estimated to affect millions of individuals worldwide (1). It is characterized by epidermal hyperplasia, altered keratinocyte differentiation, and a prominent inflammatory infiltrate dominated by T cells. Psoriasis vulgaris is the most common clinical phenotype and is recognized by the formation of scaly, thickened, erythematous plaques of varying size. The second-most common phenotype is guttate psoriasis which often occurs in younger individuals and is characterized by widespread droplet-like erythematous papules (2). While psoriasis vulgaris is recognized as a chronic remitting-relapsing condition, guttate psoriasis has more diverse disease trajectories ranging from complete remission to evolving into chronic psoriasis vulgaris (3).

Genome-wide association studies have identified the Human leukocyte antigen (HLA) allele, HLA-C*06:02, as the strongest genetic risk factor for developing psoriasis, alongside genes related to type I interferon signaling and the interleukin (IL)-23/IL17 axis (4,5). The observation of oligoclonal expanded T cells in the epidermis of resolved lesions indicates antigen-driven T cell proliferation (6–8). HLA genes are among the most polymorphic human genes (9). They encode proteins with the primary function of presenting self- and non-self-antigen derived peptides on the cell surface to be recognized by the T cell receptor (TCR) of T lymphocytes. HLA class I molecules (Classical: HLA-A, HLA-B, HLA-C; Non-classical: HLA-E, HLA-F, HLA-G) primarily present peptides between 8–12 amino acids in length, with a strong preference for nonamers (10,11). HLA class II molecules (HLA-DP, HLA-DQ, HLA-DR), on the other hand, present longer peptides - often 12–20 amino acids in length with a preference for 14–16-mers (12,13).

Over the past decade, a handful of psoriasis-associated antigens have been identified. These include peptide antigens derived from LL37, ADAMTS-like protein 5 (ADAMTSL5), keratin-17 (KRT17), serpin family B member 3/4 as well as lipid antigens presented by CD1a molecules (14–19). Psoriasis vulgaris and guttate psoriasis can both be triggered by infections caused by group A streptococci (GAS), especially in individuals carrying *HLA-C*06:02* (20,21). The phenomenon of GAS-triggered psoriasis is, among other theories, hypothesized to result from molecular mimicry between peptides derived from streptococcal antigens and human self-antigens, leading to expansion of T cell populations able to recognize and respond to common epitopes that are likely to be presented by HLA-C*06:02 in the affected skin (22,23). While these discoveries have shed light on the autoimmune nature of psoriasis, the antigenic landscape of psoriatic skin remains poorly defined.

In this study, we apply immunopeptidomics - a mass spectrometry-based strategy that directly identifies peptides bound to HLA molecules (24–26) - to profile the antigen landscape in the skin from patients with psoriasis vulgaris, guttate psoriasis and healthy individuals. This strategy allows for the systematic discovery of naturally presented self-peptides and represents a deep coverage approach to antigen exploration in psoriasis. By identifying candidate HLA class I and class II ligands identified in human lesional, non-lesional and healthy skin biopsies, our study uncovers potential antigenic triggers of T cell activation, providing a resource for mechanistic studies and advancing the search for antigen-specific immunotherapies in psoriasis.

## Methods

### Sample collection

We included patients with psoriasis (n=20) and sex- and age-matched individuals without skin disease (n=16). Patients had no topical treatment for 2 weeks and no systemic or light therapy for 4 weeks prior to biopsy collection. All participants were ≥18 years old and not pregnant. One 4 mm (13 mm^2^) or two 3 mm (2 × 7 mm^2^) lesional and non-lesional biopsies were obtained from each individual. The biopsies were snap frozen in liquid nitrogen and stored at -80□. All participants gave written informed consent. Some healthy skin samples (6 of 16) were obtained from the Australian Donation and Transplantation Biobank and collected from deceased organ donors (27). The study was conducted in accordance with the Declaration of Helsinki. The study was approved by the Ethics Committee of the Capital Region of Denmark (H-19036270), the Monash University Human Research Ethics Committee (27876), the Alfred Hospital Human Research Ethics Committee (244/23) and the Danish Data Protection Agency.

### Isolation of HLA-bound peptides from Skin biopsies

Frozen biopsies were cryogenically ground using a CryoMill (Retsch GmbH, model no. MM400) for four minutes at 30 Hz with 200 μL of lysis buffer (0.5% IGEPAL CA-630 [Sigma-Aldrich, cat. no. I8896]; 50mM Ultra-Pure Grade Tris, pH 8 [Astral Scientific, cat. no. BIO3094T]; 150mM NaCl [Supelco]; cOmplete^TM^ Protease Inhibitor Cocktail [Roche] in water) in a 5 ml stainless steel canister with a 10 mm ball that had been pre-cooled by submersion in liquid nitrogen. Cryomilled biopsy material was transferred to 2 mL Eppendorf tubes, then the CryoMill canister was washed twice with 200 μL of lysis buffer and combined with the originally transferred material. The Eppendorf tubes were rolled at 4□for 45 minutes and the resulting lysates were cleared from cell nuclei by centrifugation (15 minutes at 15,900 × g). The nuclear pellets were stored at -80 □for later DNA isolation and HLA-C*06:02 genotyping. Cleared lysates were added to 100 μl of pre-washed protein A agarose resin (Captiva®, Repligen) (precolumn) and rolled for a minimum of one hour at 4□. This precolumn resin was removed by passing the suspension through a Mobicol spin column (MoBiTech GmBH). The flow through material was incubated with protein A resin-bound HLA class I specific monoclonal antibody (W6/32, 200 ul of resin and 400 μg Ab per sample) with rolling overnight at 4□. The next day, W6/32 resin including captured HLA class I molecules was separated using Mobicol spin columns and the remaining lysate incubated with protein A agarose resin bound to a cocktail HLA class II Abs (400 μg of SPV-L3 (HLA-DQ), B7/21 (HLA-DP) and LB3.1 (HLA-DR) in a 1:1:1 ratio) with rolling for a minimum of one hour at 4□after which the Mobicol spin was repeated. The retained resin, containing captured peptide-HLA class I or class II complexes were individually washed twice with 250 mM NaCl and twice with PBS on Mobicol spin columns and then eluted by two 5 minute rounds of 200 μL of 10% acetic acid. The eluates were heated at 70□for 10 minutes (Benchmark Scientific Isoblock^TM^) and after cooling, transferred to a pre-washed 5 kDa ultrafiltration units (Ultrafree®-MC-PLHCC, Merck Millipore) for centrifugation at 15,900 × g for 50 - 120 minutes. Residual peptides were recovered by repeating the filter centrifugation step with an additional 50 μL of 10% acetic acid. The peptide containing flowthroughs were stored at -80□until further processing.

### LC-MS/MS

Peptides were dried using a vacuum centrifugation system (Labconco), reconstituted in 2% (v/v) acetonitrile (Fisher Chemical), 0.1% (v/v) formic acid (Thermo Fisher Scientific) supplemented with a mixture of 11 iRT peptides (28) (Biognosys) and loaded onto the Evotips (Evosep Biosystems) as per manufacturer instructions. The Evosep One liquid chromatography system was coupled to a hybrid ion mobility-quadrupole time of flight mass spectrometer (timsTOF Pro 2, Bruker Daltonics) and tip bound material separated using the Zoom Whisper 20 SPD method. The entire sample material was loaded onto the mass spectrometer and peptides separated using an Aurora Elite C18 Column (15 cm x 75 μm x 1.7 μm; IonOpticks). The precursor selection mode on the timsTOF Pro 2 was set to data-dependent acquisition (DDA) with Parallel Accumulation-Serial Fragmentation (PASEF) enabled. A precursor m/z range between 100-1700 Da was used with the following instrument settings; capillary voltage of 1600 V, a target intensity of 30,000 and TIMS ramp of 0.6-1.6 Vs/cm^2^ for 166 ms as described in detail elsewhere (29).

### Peptide identification

PEAKS Online 11 was used to identify peptides at 5% FDR from the spectral files matched against the human proteome (FASTA: Uniprot Human_UP5640_AND_reviewedOnly downloaded on May 5, 2024) and a contaminant database containing iRT and protein A-derived peptides. Search parameters included peptides of 6-30 amino acids in length, no enzymatic digestion, no cysteine alkylation. Parent and fragment ion tolerances were set to 20 ppm and 0.2 Da, respectively. Included variable modifications were M oxidation, N-term Acetylation and N/Q deamidation. A Peaks “DB search” was performed followed by a Peaks “PTM search” incorporating all default built-in modifications with the same mass tolerance settings.

### Data deposition

The mass spectrometry proteomics data have been deposited to the ProteomeXchange Consortium via the PRIDE partner repository with the dataset identifier PXD071219 (30).

### Bioinformatic analysis and data visualization

We retained 8-12-mers among W6/32 immunoprecipitated peptides (designated as HLA class I ligands) and 12-20-mers among SPV-L3, B7/21 and LB3.1 immunoprecipitated peptides (designated as HLA class II ligands); iRT and contaminant Protein A-derived peptides were removed from the dataset.

One set of HLA-C*06:02-positive lesional and non-lesional HLA class I data from the same patient with psoriasis vulgaris were removed due to very low yield (<600 peptides).

CSV outputs from Peaks Online were used in Perseus 2.0.5.0 to filter datasets based on peptide identity and source protein. DeepVenn (https://www.deepvenn.com) was used to generate Venn diagrams. HLA consensus motifs, GibbsCluster results and Pearson correlation between peptide clusters, reference HLA motifs and binding prediction were performed via Immunolyser 2.0 (31). Graphs were generated in GraphPad Prism (v10.4.2). Peptide binding affinities towards specific HLA allotypes were predicted using NetMHCpan 4.2 (32).

### HLA-C*06:02 genotyping

DNA was extracted from the nuclear pellets, using a DNeasy® Blood and Tissue kit (Qiagen), according to the manufacturer’s instructions for cultured cells. The extracted DNA from each sample was tested for the presence of the HLA-C*06:02 allele using a polymerase chain reaction (PCR) with the sequence-specific primers. Two distinct HLA-C target regions were tested: the forward primer 367 (F367) (5’-TAC TAC AAC CAG AGC GAG GA-3’) and reverse primer 127 (R127) (5’-GGT CGC AGC CAT ACA TCC A-3’) were used to amplify the first target region. For the second target region, forward primer 27 (F27) (5’-CCG AGT GAA CCT GCG GAA A-3’) was used with R127. The primer sequences were obtained from previously published methods, both primer sets yielded identical results (33,34). A GAPDH PCR served as a positive control: GAPDH forward primer CCA CAG TCC ATG CCA TCA C, GAPDH reverse GTT GCT GTA GCC AAA TTC GTT G. All PCR reactions (20 µL final volume) contained 10X PCR buffer (1X), 10 µM primers (0.5 µM each), 10 mM dNTPs, Q solution, and Taq DNA polymerase. HLA-C typing reactions used primer pairs F367/R127 or F27/R127 with 0.625 mM MgCl□, while GAPDH control reactions contained 2.5 mM MgCl□. The amplification consisted of an initial denaturation at 95°C for 5 minutes, followed by 30 cycles of denaturation at 96°C for 25 seconds, annealing at 55°C for 50 seconds, and extension at 72°C for 50 seconds. A final extension was performed at 72°C for 1.5 minutes, followed by a hold at 4°C.

### Immunohistochemistry

Immunohistochemical studies were performed on formalin-fixed paraffin-embedded (FFPE) sections using the fully automated Omnis instrument (Agilent/Dako, Glostrup, Denmark). Tissue sections (5 µm) were dewaxed, antigen retrieval was carried out using EnVision™ FLEX Target Retrieval Solution (TRS) High pH (Agilent/Dako, #GV804) for 24 minutes at 97°C. Endogenous peroxidase activity was blocked for 5 minutes at 32°C using Envision Peroxidase Blocking Reagent (part of the GV800 kit (Agilent/Dako)). Sections were stained for 30 minutes at 32°C with the primary antibodies: rabbit polyclonal anti-HLA-C (Thermo Fisher Scientific, MA, US; Cat. no. PA5-79367) 1:3000; mouse monoclonal anti-PI3 (Abcam, Cambridge, UK; Cat. no. ab81680, Clone TRAB2O) 1:800; mouse monoclonal anti-CD3 (Leica Biosystems, Sheffield, UK; Cat. no. NCL-L-CD3-565, Clone LN10) 1:50; rabbit monoclonal anti-CD4 (Epitomics - AH Diagnostic, Aarhus, DK; Cat. no. AC-0173, Clone EP204) 1:25; and mouse monoclonal anti-CD8 (Agilent/Dako, Glostrup, DK; Cat. no. M7103, Clone C8/144B) 1:200. Detection was performed with EnVision™ FLEX /HRP Detection Reagent (Agilent/Dako, #GV800 with Rabbit Linker #GV809) for rabbit antibodies or EnVision™ FLEX /HRP Detection Reagent (Agilent/Dako, #GV800 with Mouse Linker #GV821) for mouse antibodies. Reactions were visualized with EnVision™ Flex DAB+ Chromogen system (Agilent/Dako, #GV825) according to the manufacturer’s instructions. Finally, slides were counterstained with Hematoxylin and mounted.

### Statistical analysis

Kruskal-Wallis test followed by uncorrected Dunn’s test was used to compare peptide count distributions between groups.

## Results

To characterize the landscape of HLA class I and II-bound peptides in psoriatic versus unaffected skin, we enrolled 36 participants: 12 patients with psoriasis vulgaris, eight patients with guttate psoriasis and 16 healthy controls (Table 1). The proportion of females was higher in the guttate psoriasis group compared to the psoriasis vulgaris and healthy control groups. Disease severity ranged from mild to severe across the patient population. Patients with psoriasis vulgaris had longer mean disease duration compared to patients with guttate psoriasis (14.9 versus 8.1 years). Local disease activity, measured by local Psoriasis Severity Index (PSI) at the biopsy site was higher in the psoriasis vulgaris group compared to the guttate psoriasis group (5.6 and 3.9, respectively). Four out of eight patients with guttate psoriasis and four out of 12 patients with psoriasis vulgaris were HLA-Cw6 carriers.

**Table 1-.**
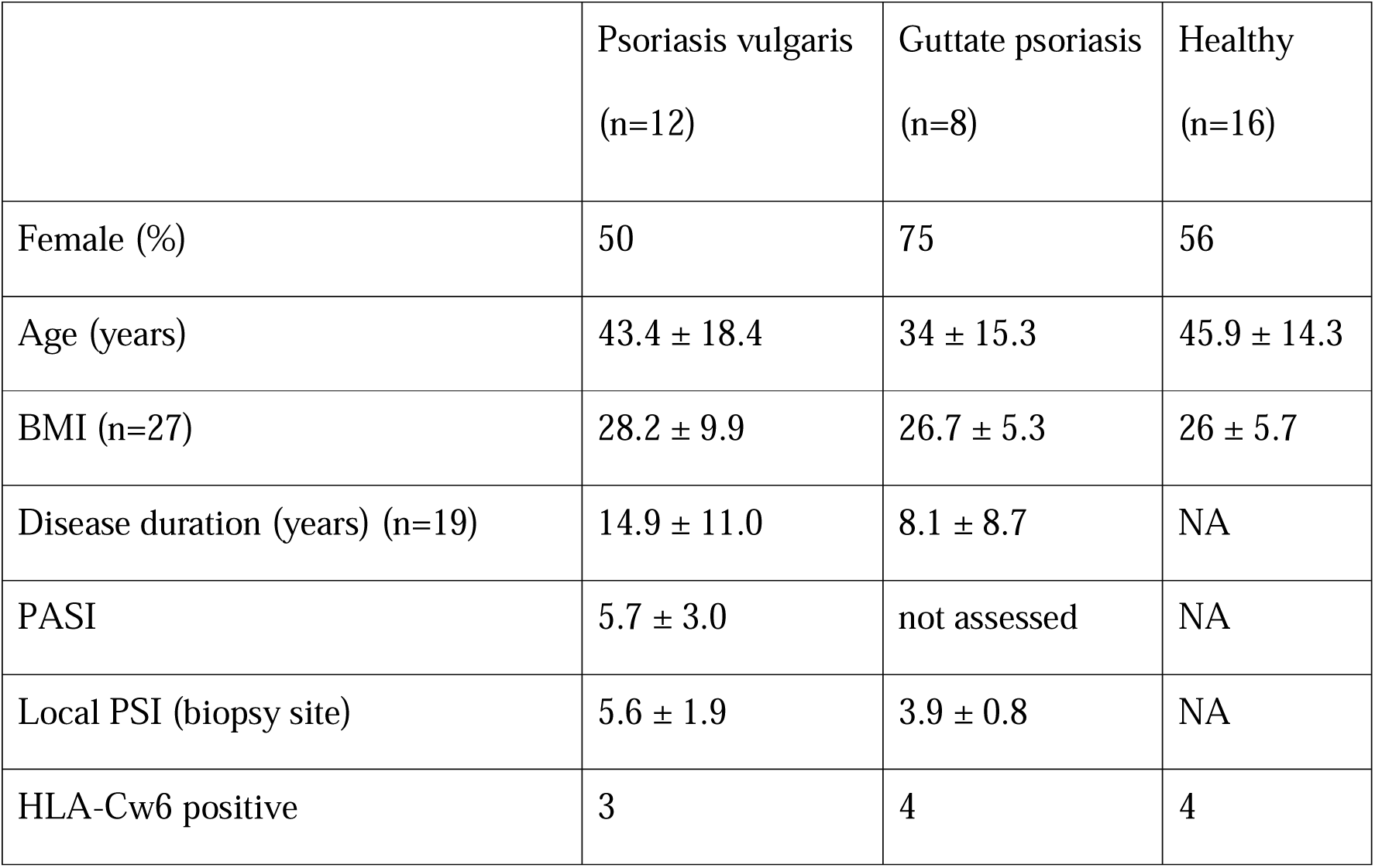
demographic and clinical information. Continuous variables are presented as mean ± SD; categorical variables as n or percentage

|  | Psoriasis vulgaris<br>(n=12) | Guttate psoriasis<br>(n=8) | Healthy<br>(n=16) |
| --- | --- | --- | --- |
| Female (%) | 50 | 75 | 56 |
| Age (years) | 43.4 ± 18.4 | 34 ± 15.3 | 45.9 ± 14.3 |
| BMI (n=27) | 28.2 ± 9.9 | 26.7 ± 5.3 | 26 ± 5.7 |
| Disease duration (years) (n=19) | 14.9 ± 11.0 | 8.1 ± 8.7 | NA |
| PASI | 5.7 ± 3.0 | not assessed | NA |
| Local PSI (biopsy site) | 5.6 ± 1.9 | 3.9 ± 0.8 | NA |
| HLA-Cw6 positive | 3 | 4 | 4 |

### A higher number of immunopeptides was identified in lesional psoriasis samples

We purified HLA-bound peptides from all skin biopsies via serial immunoprecipitation of HLA class I and class II and analysed the bound peptide ligands using mass spectrometry. The analyses yielded a total of 99,204 HLA class I- and II-bound (6-30-mer, PTMs removed, non-redundant) peptides identified across all samples (Supplementary Data 1).

We filtered our analysis for the predominant peptide lengths: 8-12-mer peptides for HLA class I eluates and 12-20-mer peptides for HLA class II eluates. This retained 54,816 HLA class I ligands and 28,139 HLA class II ligands.

We found elevated numbers of uniquely identified peptides isolated from lesional skin. On average, we identified 6,201 HLA class I ligands in lesional psoriasis vulgaris, 4,798 in lesional guttate psoriasis and 2,359 in unaffected skin with minimal variation between non-lesional samples from patients and skin from healthy donors (Fig. 1A). Within each lesional/non-lesional sample pair, an average of 60.6% (SD 13.5) and 63.1% (SD 13.6) of the HLA class I-bound peptides were exclusively identified in the lesional sample from patients with psoriasis vulgaris and guttate psoriasis, respectively (Fig. 1A). Lower numbers of HLA class II ligands were identified compared with HLA class I ligands. Mean peptide counts per sample were 3,103 and 2,213 in lesional psoriasis vulgaris and guttate psoriasis, respectively, and 1,026 in non-lesional and healthy skin (Fig. 1B). This is in line with HLA class II expression predominantly found on professional APCs that constitute a minority of the cells present in skin, while HLA class I is expressed by nearly all cells. Similar to the HLA class I findings, the majority of HLA class II ligands were identified exclusively in the lesional biopsies of both psoriasis vulgaris (67.4%, SD 12.2) and guttate psoriasis donors (60.0%, SD 14.8) (Fig. 1B).

**Figure 1.**
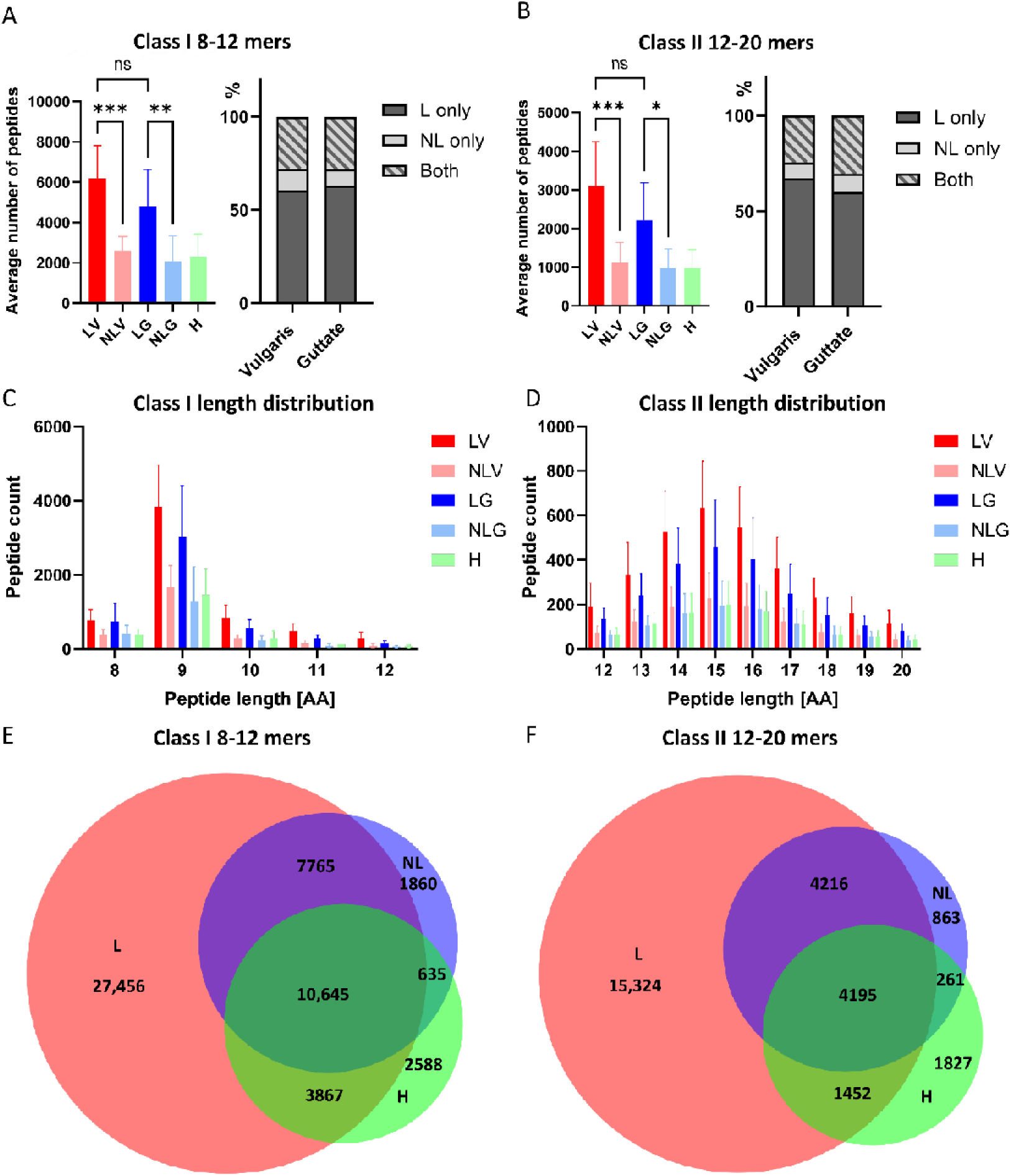
Characteristics of HLA ligands isolated from psoriatic and unaffected skin. **(A-B)** The average numbers and overlap of unique HLA class I- and II-bound peptides eluted using the pan-HLA class I antibody W6/32 or the HLA class II antibodies SPV-L3, B7/21 and LB3.1. Error bars represent standard deviation. **(C-D)** Relative length distribution of peptides bound to HLA class I and II, respectively (HLA class I) or 15-mers (HLA class II). Error bars represent standard deviations. (E-F) Venn diagrams of all HLA class I- and II-bound peptides identified in lesional, non-lesional and healthy samples. LV = lesional psoriasis vulgaris (HLA class I: n = 11, HLA class II: n = 12), NLV = non-lesional psoriasis vulgaris (HLA class I: n = 11, HLA class II: n = 12), LG = lesional guttate psoriasis (n = 8), NLG non-lesional guttate psoriasis (n = 8), H = healthy (n = 16), L = lesional (HLA class I: n = 19, HLA class II: n = 20), NL = non-lesional (HLA class I: n = 19, HLA class II: n =20), AA = amino acids. *: p < 0.05, **: p < 0.01, ***: p < 0.001 (Kruskal-Wallis test).

Global peptide length distribution analysis confirmed expected HLA class I and class II peptide length preferences across all sample groups, with 9-mer peptides dominating HLA class I ligands and 14-16-mer peptides most prevalent among the HLA class II ligands (Fig. 1C-D).

We next investigated the extent of peptide overlap across all lesional, non-lesional and healthy samples combined. Peptides detected in non-lesional and healthy samples were mostly also represented in lesional samples (87.5% and 79.2%, respectively). On the other hand, 27,457 HLA class I-bound peptides (45%) and 15,325 HLA class II-bound peptides (54%) were exclusively detected in lesional skin (Fig. 1E-F).

### The HLA class I immunopeptidome shares specific peptides in lesional skin from C*06:02-positive patients

To further investigate the HLA I immunopeptidome, we first used GibbsCluster 2.0 to identify peptide groups and classify the peptides into potential HLA I allotype specific ligands based on their sequence motifs (35,36). In six of seven patients genotyped as HLA-C*06:02-positive, we were able to identify a cluster with a preferred presentation of arginine residues at position P2/7 and hydrophobic amino acids at the C-terminal (PΩ) position, characteristic of HLA-C*06:02 ligands (Fig 2A). HLA-C*06:02 is often co-inherited together with HLA-B*57:01 and we could also identify a likely HLA-B*57:01 cluster in five of seven HLA-C*06:02-positive patients with an S/T/A preference on P2 and W/F preference on PΩ (Fig 2A).

**Figure 2.**
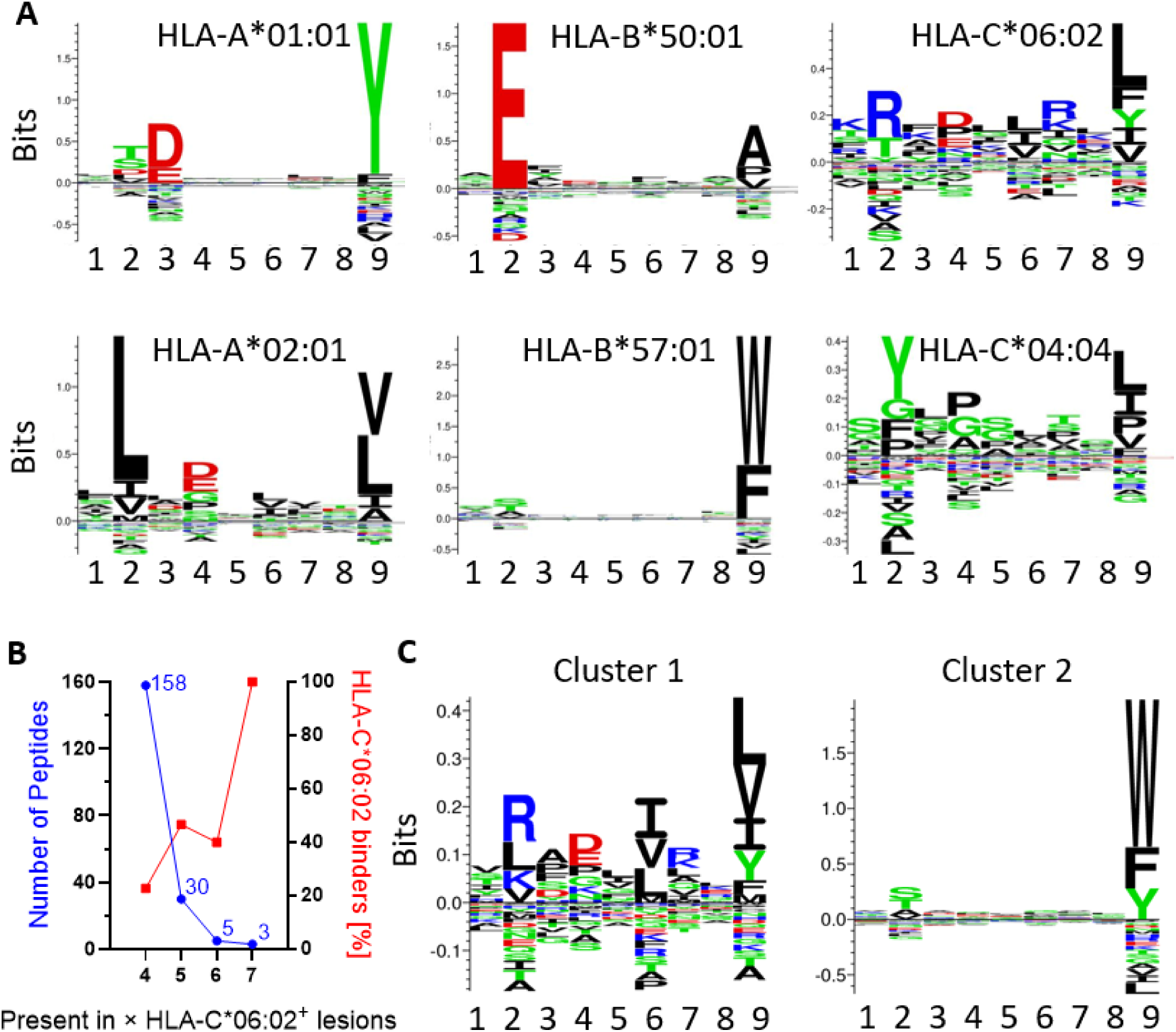
Peptides repeatedly detected in lesions of HLA-C*06:02-positive patients align well with the HLA-C*06:02 reference motif. **(A)** GibbsCluster groups with the highest KLD scoring result for 6 clusters of a representative lesional sample from an HLA-C*06:02-positive individual is able to detect distinct sequence motifs that can be readily attributed to putative HLA class I allotypes for this donor. Prediction of source HLA with Immunolyser 2.0. **(B)** Peptides commonly detected in four to seven of the HLA-C*06:02-positive lesions. Plotted on the left y-axis in blue is the number of peptides identified in increasing numbers of HLA-C*06:02-positive lesional samples but not detected in any of the non-lesional or healthy tissues. The percentage of these peptides predicted to be HLA-C*06:02 binders is plotted on the right y-axis in red. **(C)** GibbsCluster grouping of 196 peptides from (B) found in 4-7 lesions of HLA-C*06:02-positive patients align with HLA-C*06:02-like (cluster 1) and HLA-B*57:01-like (cluster 2) motifs.

Next, to identify potential pathogenic peptides originating from the major risk gene HLA-C*06:02, we retained only HLA class I ligands consistently detected in lesional skin from HLA-C*06:02-positive patients and absent from all non-lesional and healthy skin samples, irrespective of HLA composition (Fig. 2B, Supplementary table 1).

The three nonamers, MRASSFLIV from peptidase inhibitor 3 (PI3), SNSDVIRQV derived from ubiquitin carboxyl-terminal hydrolase enzyme L5 (UCHL5) and FRSEDIKRL from conserved oligomeric Golgi complex subunit 4 (COG4) were present in all seven lesions of HLA-C*06:02-positive patients and were not identified in unaffected skin. They were all predicted to bind strongly to HLA-C*06:02 (MRASSFLIV: %Rank = 0.0932; SNSDVIRQV: %Rank = 0.1437; FRSEDIKRL: %Rank = 0.0049).

Five additional peptides were identified in six of the seven lesions from HLA-C*06:02-positive patients including KVVDVVRNL derived from interferon-induced GTP binding protein Mx1 (MX1) and TRTDKVRAL from protein transport protein Sec61 subunit alpha which were also predicted to bind to HLA-C*06:02 (KVVDVVRNL: %Rank = 0.0764; TRTDKVRAL: %Rank = 0.0047). The 8-mer NTPFLNID from keratinocyte differentiation-associated protein (KRTDAP), the 10-mer VVDPFSKKDW and the 11-mer KDWQRRVATWF from ribosomal subunit proteins present in six lesions from HLA-C*06:02-positive patients were not predicted HLA-C*06:02-binders. A follow-up GibbsCluster analysis of the top 196 peptides detected in at least four of seven HLA-C*06:02^+^ lesions and not detected in non-lesional skin segregated the peptide sequences with binding characteristics of HLA-C*06:02 or HLA-B*57:01 ligands (Fig. 2C).

### PI3, the source protein of MRASSFLIV, is upregulated in lesional psoriatic epidermis

The consistent detection of MRASSFLIV in all HLA-C*06:02-positive patients and its exclusive detection in lesions warranted further investigation. The protein source, PI3, has well-established associations with psoriasis and we sought to examine whether PI3 was expressed in regions exhibiting HLA-C expression and cytotoxic T-cell infiltration. Immunohistochemical staining revealed high expression of PI3 in psoriatic lesions, predominantly localized to the epidermis, the same region where HLA-C is primarily expressed and where CD8^+^ T cells accumulate in psoriatic lesions (Fig. 3A-F).

**Figure 3.**
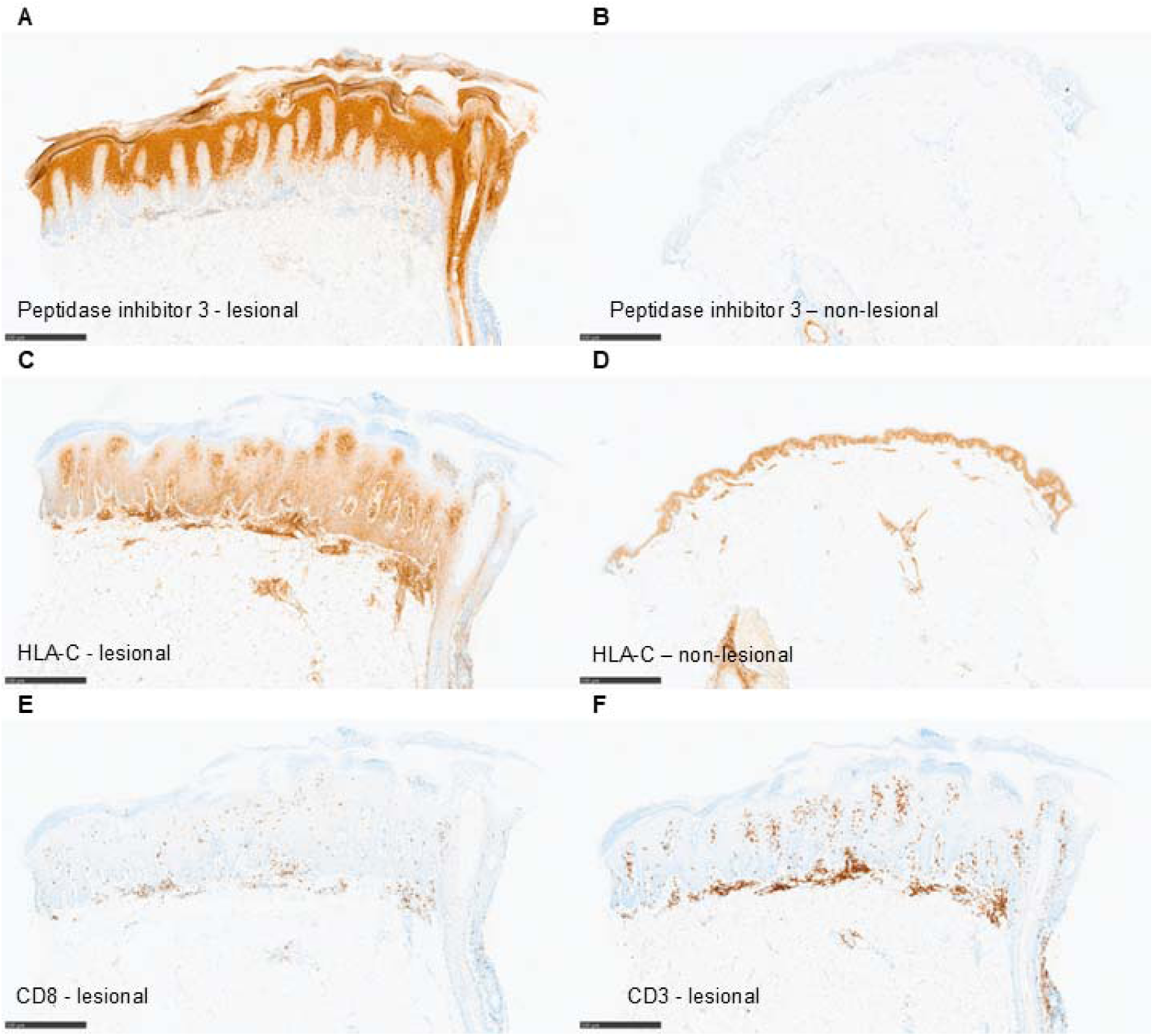
Immunohistochemistry indicates strong PI3 expression in lesional epidermis alongside HLA-C expression and cytotoxic T cell infiltration. Immunohistochemical staining of peptidase inhibitor 3 (PI3) **(A, B)** and HLA-C **(C, D)** expression in lesional versus non-lesional skin and CD8^+^ **(E)** and CD3^+^ **(F)** cells in lesional skin. The immunohistochemical staining data **(A-F)** stem from the same individual and serve as a representative example of four patients. The scale bars represent 500 μm.

### KRTDAP contributes multiple lesion-specific HLA class II ligands

Given the clonal expansion of CD4^+^ Th17 cells in psoriasis we also investigated the landscape of HLA class II ligands in psoriatic skin (37). HLA class II restriction of peptides can be considered more promiscuous than that observed with HLA class I due to the ability the typically longer peptide ligands to contain several HLA class II allotype specific binding cores. Six HLA class II-bound peptides were exclusively identified in lesions and were present in at least half of the lesional biopsies analysed. Three of these six peptides were derived from KRTDAP, a relatively short protein of 99 amino acids that is highly enriched in skin. One of the lesion-specific KRTDAP-derived peptides, RPEAF<u>NTPFLNID</u>K, notably overlaps with the octamer peptide previously highlighted among peptides exclusively identified in lesional biopsies and present in all HLA-C*06:02 positive samples (underlined). Taken together, the three KRTDAP-derived peptides were identified in 9/12 psoriasis vulgaris and 5/8 guttate psoriasis patients. We investigated whether the repeated lesional appearance of peptides from KRTDAP was a consequence of this antigen generally dominating the HLA class II immunopeptidome in skin, however we found that it only contributed to 0.6% of all peptides. In comparison, the proteins that contributed the highest number of HLA class II bound peptides were the abundant serum proteins apolipoprotein B-100 (1.9%) and complement C3 (1.4%).

### Global source protein-level analysis reveals the representation of several candidate autoantigens in the immunopeptidome

The HLA haplotype diversity of the study participants was reflected in differences in peptide antigen presentation and sequences consistent with corresponding HLA-allotype-specific binding motifs (Fig. 2A). To be able to make comparisons between patients regardless of their HLA typing, we sought to determine whether the source proteins of peptides processed and presented in lesional tissues were consistent across patients. We first filtered for peptides that were unambiguously assigned to have originated from a single source protein (∼90% of all peptides) to then identify proteins processed and presented in detectable amounts exclusively in lesions. From a total of 9,796 unique source proteins presented by HLA class I molecules, 2,673 were exclusively detected in lesional skin from patients with psoriasis vulgaris or guttate psoriasis (Fig. 4A). Fewer source proteins were detectable in the HLA class II eluates amounting to a total of 2,717 of which 935 were exclusively detected in lesions (Fig. 4B). Among lesion-specific source proteins, 10 were detected in at least half of the HLA class I samples, and 5 were detected in at least half of the HLA class II samples (Fig. 4C-D). Supplementary Table 2 provides an overview of these proteins including the number of identified peptide sequences, presentation on either HLA class, HLA binding properties, skin layer-specific expression in psoriasis, general protein tissue expression and protein function. Overall, all the source proteins in Supplementary Table 2 are known to be expressed in skin. HLA class I ligand sources were annotated to be intracellular or membrane-bound proteins while HLA class II ligand sources were all from known or predicted secreted proteins. An exception was the secreted protein PI3, found in the HLA class I immunopeptidome. Further investigation of the identified PI3 peptides revealed that they all originated from either the signal peptide (which is cleaved intracellularly) or the pro-peptide and the most frequently observed peptide was the aforementioned N-terminal signal peptide (MRASSFLIV). The number of unique peptide sequences from each protein varied from three to 34 except for DUSP2 with only one peptide. Functional annotations revealed a broad distribution across biological processes. These included proteins involved in RNA processing, DNA repair, lipid and amino acid metabolism, (de-)phosphorylation, immunomodulation, intracellular homeostasis and extracellular structure. No direct protein-protein interactions were identified among the 19 source proteins based on the interaction databases available via the STRING database (38).

**Figure 4.**
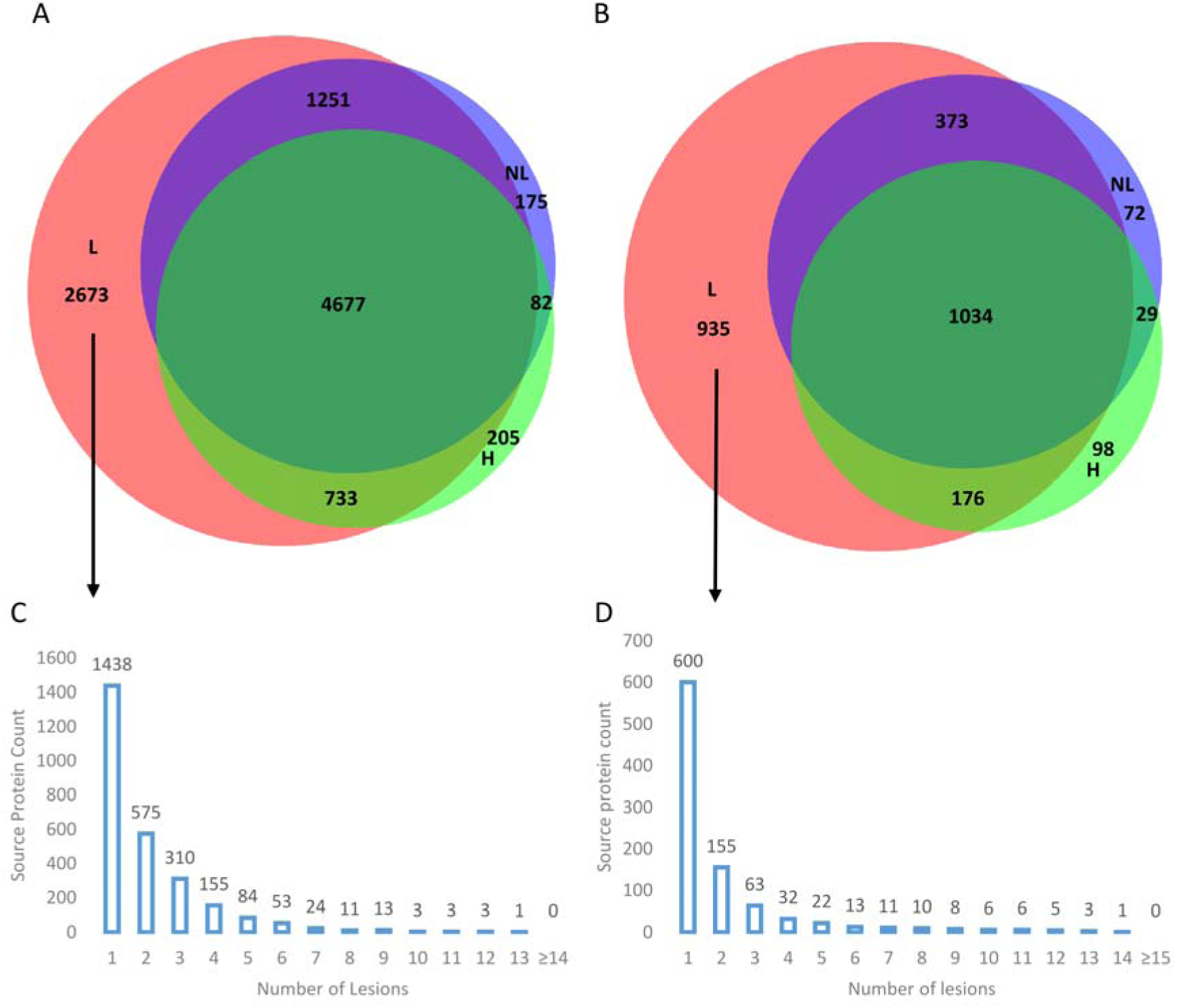
Source protein overlap between lesional and unaffected skin and analysis of lesion-specific source proteins. **(A-B)** Venn diagrams of source protein identities unambiguously assigned based on HLA class I and class II ligands. **(C-D)** Source protein prevalence for HLA class I and HLA class II source proteins which were only identified in lesions (zero values not shown).

Additionally, we explored whether the presence of specific source proteins could differentiate between psoriasis vulgaris and guttate psoriasis or correlated with *HLA-Cw6* carrier status (specifically for HLA class I-bound peptides). However, no individual source protein was found to discriminate between these psoriasis subtypes or the *HLA-Cw6* status.

## Discussion

In this study, we used immunopeptidomics to map the HLA class I and class II–bound peptide repertoires of lesional and non□lesional skin from patients with psoriasis and from healthy individuals. Our workflow, leveraging serial pan□HLA class I and class II immunoprecipitations and high□resolution mass spectrometry, enabled the identification of over 99,000 distinct peptides from skin punch biopsies, underscoring the capacity of this approach to capture the *in vivo* antigenic landscape. With this approach, we have identified several candidate autoantigens at the peptide level for HLA-C*06:02^+^ individuals and source protein level for all patients based on their representation in the immunopeptidomes from the HLA class I and II antigen presentation pathways.

Consistent with the heightened inflammatory milieu and increased thickness of active plaques, lesional skin yielded substantially higher numbers of identified immunopeptides compared to unaffected skin. This likely reflects increased keratinocyte turnover, immune cell infiltration, and cytokine□driven upregulation of the antigen processing machinery (39,40). The largest differences in peptide composition between lesional and unaffected skin was observed for HLA class II bound peptides, in line with the presence of infiltrating antigen□presenting cells under inflammatory stimuli (41).

Notably, analysis of HLA-C*06:02 positive samples revealed peptides from PI3 and MX1 predicted to bind strongly to HLA-C*06:02 that were identified in six or all seven HLA-C*06:02 positive lesional samples, and other peptide sequences from PI3 and MX1 were detected in HLA-Cw6-negative lesional samples. In particular, the PI3–derived N-terminal signal peptide fragment, MRASSFLIV, represents a candidate for further validation of its immunogenicity in individuals with psoriasis (42). Interestingly, PI3 was initially discovered in psoriatic skin and has previously been suggested as a biomarker correlating with psoriasis severity (43–45). Furthermore, PI3 expression has been demonstrated to be induced by known triggers of psoriasis in HLA-C*06:02-positive individuals such as skin trauma, infections and, particularly, streptococcal soft tissue infection (46–49). Additionally, the MX1-derived peptide KVVDVVRNL was detected among HLA class I ligands in six of seven lesional samples from HLA-C*06:02 positive individuals. As a type I interferon-induced protein previously demonstrated to be upregulated in lesional psoriatic skin, MX1 is well situated to be a source of pathogenic antigens both in the initiation and sustained disease phases (50–52). Both of the highlighted peptides consist of an arginine on position two or seven, thereby aligning well with the HLA-C*06:02 motif (53–56) and were also predicted HLA-C*06:02 binders via NetMHCpan 4.2 (32). Further investigation of their chemical interactions with pathogenic psoriatic T cell receptors could provide mechanistic insight into their potential involvement in the disease pathogenesis (57).

In the HLA class II compartment, KRTDAP stood out as a skin tissue enriched source protein, represented by multiple immunopeptides exclusively detected in psoriatic plaques and notably one of the peptides (RPEAFNTPFLNIDK) contained a sequence (NTPFLNID) also frequently represented in HLA-C*06:02 positive lesions. The expression of KRTDAP itself has, to our knowledge, not been reported to be altered in psoriatic skin. However, it is part of the epidermal differentiation complex in the PSORS4 locus and we have previously identified increased levels of KRTDAP in the plasma of patients with psoriasis (58). Whether the presentation of KRTDAP fragments to both CD4□and CD8+ T cells contributes to amplification of local and systemic inflammation will be an interesting avenue for future research.

Recently, SerpinB3/4 immunopeptides have been suggested as autoantigens, especially in eczematized psoriasis, indeed we could also confirm 27 of these previously reported immunopeptides and report further peptides from various Serpins (18). In contrast, for previously identified psoriasis autoantigens LL37 and ADAMTSL5 with reported T cell reactivity, we did not detect any immunopeptidomes. Our unbiased immunopeptidomics approach identified several alternative autoantigenic peptides whose processing and presentation are tightly linked to known disease triggers (59,60). Thus, the results of this study provide a resource for future investigation of potentially pathogenic peptide antigens in psoriasis. The presence of an antigen, while a prerequisite for eliciting an immune reaction, does not inform about its immunogenicity (61). Hence, an important next step will be to evaluate whether the proposed candidate autoantigens are recognized by and capable of activating T cells from patients with psoriasis.

Several methodological limitations warrant consideration. First, differences in expressed HLA alleles and inter-sample variability in peptide yield between lesional and non-lesional tissues, influenced by factors such as limited biopsy size, variable inflammation intensity and the DDA mass spectrometry method introduce challenges for quantitative comparisons (62). Second, the isolation of pan-HLA I and pan-HLA II peptides precludes definitive assignment of individual peptides to specific HLA allotypes - future studies employing allele-specific immunoprecipitation, especially of HLA-C, will be necessary to resolve this limitation.

In conclusion, we individually characterised the HLA class I and HLA class II immunopeptidomes of the skin from patients with psoriasis vulgaris and guttate psoriasis, identifying shared HLA class I-bound peptides presented by HLA-C*06:02-positive individuals that were identified exclusively in lesional skin. Our comprehensive immunopeptidomic profiling of psoriatic skin highlights promising candidate autoantigens such as PI3, MX1 and KRTDAP as prominent sources of lesion-specific HLA□presented peptides. Our findings offer a valuable framework for future studies aiming to elucidate antigen□specific mechanisms in psoriasis.

## ACKNOWLEDGEMENTS

We would like to thank all patients and healthy volunteers for their participation in the study. We also gratefully acknowledge the generosity of the deceased organ donors and their families in providing valuable tissue samples to advance medical research. The Australian Donation and Transplantation Biobank is supported by the Department of Infectious Diseases and Immunology and the Australian Centre for Transplant Excellence and Research (ACTER) at Austin Health. The authors thank Kirti Pandey for the expert care and maintenance of the timsTOF Pro 2 mass spectrometer and Rochelle Ayala for outstanding lab management.

## ABBREVIATIONS

AA: Amino Acids
ADAMTSL5: ADAMTS-like Protein 5
APC: Antigen-Presenting Cell
BMI: Body Mass Index
CD1a: Cluster of Differentiation 1a
COG4: Conserved Oligomeric Golgi Complex Subunit 4
CV: Column volume
DAB: 3,3′-Diaminobenzidine (chromogen)
DDA: Data-Dependent Acquisition
DNA: Deoxyribonucleic Acid
ERAP1: Endoplasmic Reticulum Aminopeptidase 1
FDR: False Discovery Rate
FFPE: Formalin fixed, paraffin-embedded
GAS: Group A Streptococcus
GAPDH: Glyceraldehyde-3-Phosphate Dehydrogenase
H: Healthy
HRP: Horseradish Peroxidase
HLA: Human Leukocyte Antigen
IL: Interleukin
iRT: Indexed Retention Time (peptides)
KIR: Killer-cell Immunoglobulin-like Receptor
KLD: Kullbach Leibler Distance
KRT17: Keratin 17
KRTDAP: Keratinocyte Differentiation-Associated Protein
LC-MS/MS: Liquid Chromatography–Tandem Mass Spectrometry
L: Lesional
LG: Lesional Guttate Psoriasis
LV: Lesional Psoriasis Vulgaris
MgCl□: Magnesium Chloride
MX1: Interferon-induced GTP-binding Protein Mx1
NA: Not Assessed / Not Applicable
NaCl: Sodium Chloride
NL: Non-lesional
NLG: Non-lesional Guttate Psoriasis
NLV: Non-lesional Psoriasis Vulgaris
NK: Natural Killer (cell)
PASI: Psoriasis Area and Severity Index
PBS: Phosphate-Buffered Saline
PCR: Polymerase Chain Reaction
PI3: Peptidase Inhibitor 3
PSI: Psoriasis Severity Index
PTM: Post-Translational Modification
Sec61: Protein Transport Protein Sec61 Subunit Alpha
SD: Standard Deviation
TBE: Tris-Borate-EDTA (buffer)
TCR: T Cell Receptor
UCHL5: Ubiquitin Carboxyl-terminal Hydrolase L5

